# Patient-derived parathyroid organoids as tracer and drug-screening application model

**DOI:** 10.1101/2022.03.24.485627

**Authors:** Milou E. Noltes, Luc H.J. Sondorp, Laura Kracht, Inês F. Antunes, René Wardenaar, Wendy Kelder, Annelies Kemper, Wiktor Szymanski, Wouter T. Zandee, Liesbeth Jansen, Adrienne H. Brouwers, Robert P. Coppes, Schelto Kruijff

## Abstract

Parathyroid diseases are characterized by dysregulation of calcium homeostasis and alterations in parathyroid hormone (PTH) excretion. The understanding of parathyroid hyperplastic growth and the development of parathyroid-targeted treatment and imaging tracers could benefit from *in vitro* models. Therefore, we aim to establish stem cell-derived, three-dimensional organoids representing human parathyroid tissue *in vitro*. Patient-derived hyperplastic parathyroid tissue was dispersed and parathyroid organoids (PTO) were cultured and characterized. PTO-derived cells were shown to exhibit *in vitro* self-renewal over several passages, indicative of the presence of putative stem cells. Immunofluorescence and RNA-sequencing confirm that PTO phenocopy hyperplastic parathyroid tissue. Exposure of PTO to increasing calcium concentrations and to PTH-lowering drugs resulted in a significantly reduced PTH excretion. Next to this, the PTO showed specific binding of ^11^C-methionine to the targeted receptor. Additionally, when organoids were incubated with ^99m^Tc-sestamibi, we observed a higher uptake in PTOs from patients with a ^99m^Tc-sestamibi positive scan compared to patients with a negative scan. These data show functionality of PTOs resembling the parathyroid. In conclusion, we present a patient-derived PTO culture, that recapitulates the originating tissue on gene and protein expression and functionality. This PTO model paves the way for future physiology studies and therapeutic target and tracer discovery.

## 1. Introduction

The parathyroid glands usually are four delicate structures of 3-4 mm in size, situated close to the thyroid gland, and functionally maintaining calcium homeostasis by secreting parathyroid hormone (PTH). Calcium is an essential element for the nervous, muscular and skeletal systems. Parathyroid glands consist of chief- and oxyphil cells, the former of which express calcium-sensing receptor (CaSR), and regulate the blood calcium concentration by mediating PTH secretion (1). Oxyphil cells are less prevalent than chief cells and increase in number with age (2). Parathyroid diseases are heterogeneous conditions that generally are caused by alterations in PTH secretion, resulting in dysregulation of calcium homeostasis. In primary hyperparathyroidism, a common parathyroid disease, reduced CaSR expression has been noted (3–5), resulting in uncontrolled continuous release of PTH by parathyroid hyperplasia and leading to subsequent hypercalcemia (6).

The understanding of parathyroid hyperplastic growth and the development of effective parathyroid tracers for diagnostic imaging and drugs for targeted treatment could benefit from *in vitro* models derived from human hyperplastic parathyroid tissue. To develop a model for studying the functional and proliferative properties of the parathyroid gland, two rat-derived parathyroid clonal epithelial cell lines were previously developed illustrating *in vitro* calcium–dependent PTH responses (7,8). Only one of these cell lines expressed a wide range of parathyroid-related genes (7). In further attempts to develop an *in vitro* parathyroid model, bovine parathyroid glands were cultured that showed long-term serial propagation of parathyroid cells (9–12). However, these cultures were swiftly overgrown by fibroblastic cells and rapidly lost calcium responsiveness (11,12). Besides the use of animal derived parathyroid cells, human parathyroid cell cultures have been investigated. These cultures had a low-proliferative activity and were readily terminally differentiated in a 2D environment (13–16), making them less useful for an *in vitro* model.

Organoids are 3D structures that closely recapitulate tissue architecture and cellular composition and are developed from stem cells (17). Organoid models have been developed from multiple exocrine and endocrine glandular tissues (18–26). These models have proven very useful for studying tumor behaviour and assessing drug responses, and have provided a platform for long-term *in vitro* experimentation (18–23). Hence, this study aimed to isolate human parathyroid stem cells from primary tissue and study their potential to expand and form parathyroid organoids (PTO) *in vitro* that recapitulate functional parathyroid tissue. In addition, we assess the potential of such a patient-derived PTO model to perform physiology studies and the ability to be used for the development of potential novel therapeutic targets, drug screening and imaging tracer testing.

## 2. Results

### 2.1 In vitro self-renewal and organoid formation from human parathyroid stem cells

In total, fifty hyperplastic parathyroid gland biopsies were dissociated into single cells and cell clumps using mechanical and enzymatic digestion. These cells were cultured in parathyroid medium, and resulted in the formation of parathyroid spheres (Figure 1A-B, passage 0). To determine the presence of potential stem cells in parathyroid tissue, the *in vitro* self-renewal potential of cells derived from parathyrospheres was assessed. Seventeen of 50 cultures were used to determine organoid forming efficiency, with 100% of cultures reaching passage 1 (p1), 41% reaching p2, 35% reaching p3 and 18% reaching p4 with a maximum organoid-forming efficiency (OFE) of 6.6% in p1, 2.7% in p2 and 2.9% in p3 (Figure 1B-C). Thus, self-renewal of parathyroid cells was shown. The parathyroid organoids (PTOs) continued to secrete PTH during culture (Figure 1D). To further characterize PTOs, gene expression of parathyroid markers was evaluated by quantitative Polymerase Chain Reaction (qPCR). Gene-expression analysis of parathyroid markers PTH, CaSR, and GATA Binding Protein 3 (GATA3), a transcription factor involved in the development of parathyroid tissue and in adult parathyroid cell proliferation (27), showed a stable expression between passages 1-3 (no significant differences in expression between passages, all p-values > 0.065) (Supplemental Figure 1).

**Figure 1.**
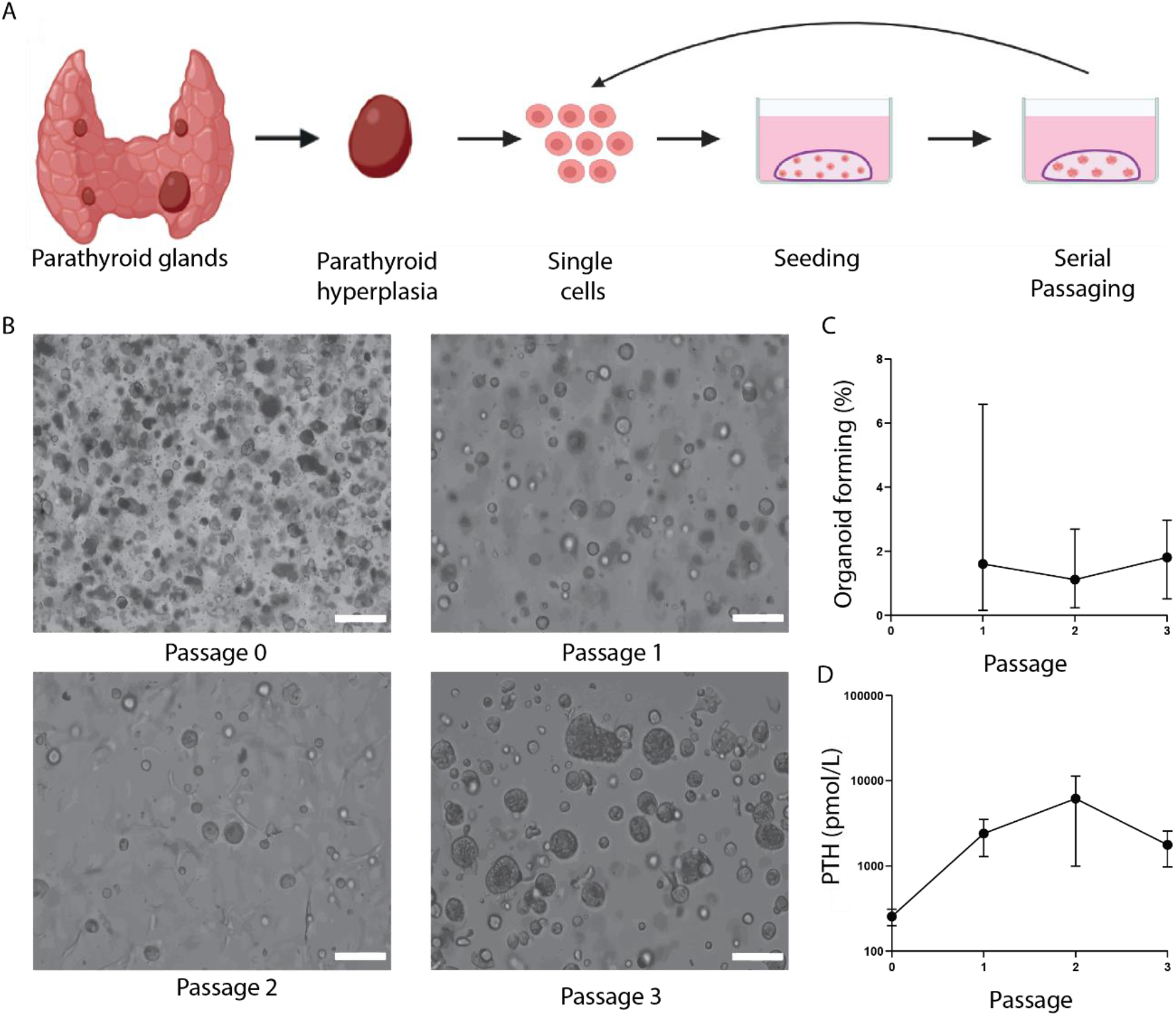
*In vitro* self-renewal of organoid-forming cells from human putative parathyroid stem cells. **A**. Schematic representation of parathyroid tissue isolation, primary tissue culture, and the self-renewal assay. **B**. Primary parathyroid spheres seven days in culture (passage 0), and PTOs at the end of passage 1, 2 and 3. Scale bar = 100 μm **C**. Self-renewal potential of PTOs during passaging (n=19 patients error bars represent range). **D.** Secreted PTH levels from PTOs during passaging (n=4 patients, error bars represent SEM). *PTO*= parathyroid organoid.

### 2.2 Characterization of parathyroid organoids

To evaluate whether the putatative parathyroid stem cells are able to differentiate into parathyroid gland specific cells, the cells were subjected to a modified Bingham protocol (28,29) involving continuously exposure to 100 ng/mL Activin, 100 ng/mL Sonic hedgehog (Shh) and 5% fetal bovine serum (FBS) for 21 days with media changes every seven days. Previous studies showed that the Bingham protocol successfully replicated parathyroid differentiation using human embryonic stem cells or tonsil-derived mesenchymal stem cells. These cells induced a significant release of PTH with response to increasing or decreasing calcium levels (28–30).

During differentiation, the cells increased slightly in size, however no further morphological changes were observed (Figure 2B). PTH secretion increased from the first week of differentiation until the second week (Figure 2C). Only in week 2 of differentiation, the PTH levels differed significantly from the PTH levels secreted prior to differentiation (average increase in PTH of 46.2% ± 21.6% in week 1 (p=0.166) and of 58.6% ± 2.6% in week 2 (p=0.002)).

**Figure 2.**
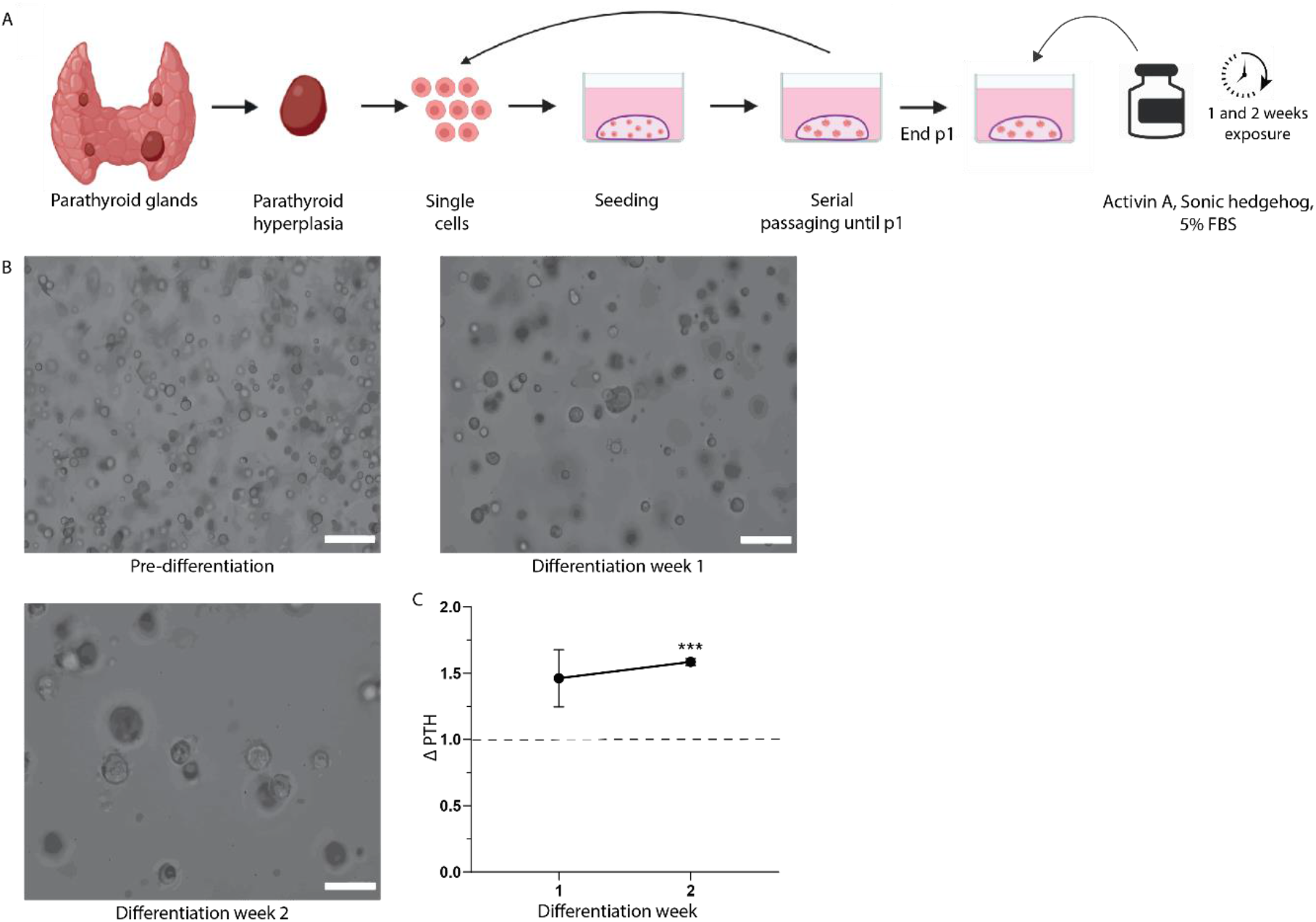
Differentiated parathyroid organoids. **A.** Schematic representation of parathyroid tissue isolation, primary tissue culture, and differentiation. Scale bar = 100 μm. **B.** PTOs before differentiation (end passage 1) and, after 1 week and 2 weeks of differentiation. **C.** Secreted PTH levels after 1 week and 2 weeks of differentiation normalized to PTH levels prior to differentiation (n=3 patients, error bars represent SEM). ***= significant (p<0.05). *PTO=* parathyroid organoid.

To further characterize PTOs and differentiated PTOs (dPTO), gene expression levels were assessed using bulk RNA-sequencing to compare hyperplastic parathyroid tissue (n=2), PTOs (n=3) and Bingham Protocol dPTOs (n=3). When assessing the number of differentially expressed genes (DEGs) between hyperplastic parathyroid tissue and PTOs, we only observed 25 up-, and 61 down-regulated genes out of 14,900 evaluated genes. Moreover, between tissue and dPTOs, we observed 50 up-, and 82 down-regulated genes out of 14,900 evaluated genes. Only two out of 35 genes of the parathyroid markers (Figure 3) were differentially expressed in both PTOs and dPTOs compared to tissue. These two genes were PTH and CHGA, both excreted factors, and both downregulated in PTO and dPTOs compared to tissue. Thus, similarity in gene expression between tissue and PTOs and dPTOs was observed, especially in parathyroid-specific genes (Figure 3). However, we did observe inter-patient variability in the gene expression of tissue (i.e. between tissue of patient 1 and 2), which was also reflected in the respective PTOs and dPTOs (Supplemental Figure 2, Supplemental Table 1). When comparing tissue, PTOs and dPTOs of one single patient (e.g. of patient 1), limited variability was observed (Supplemental Figure 2).

**Figure 3.**
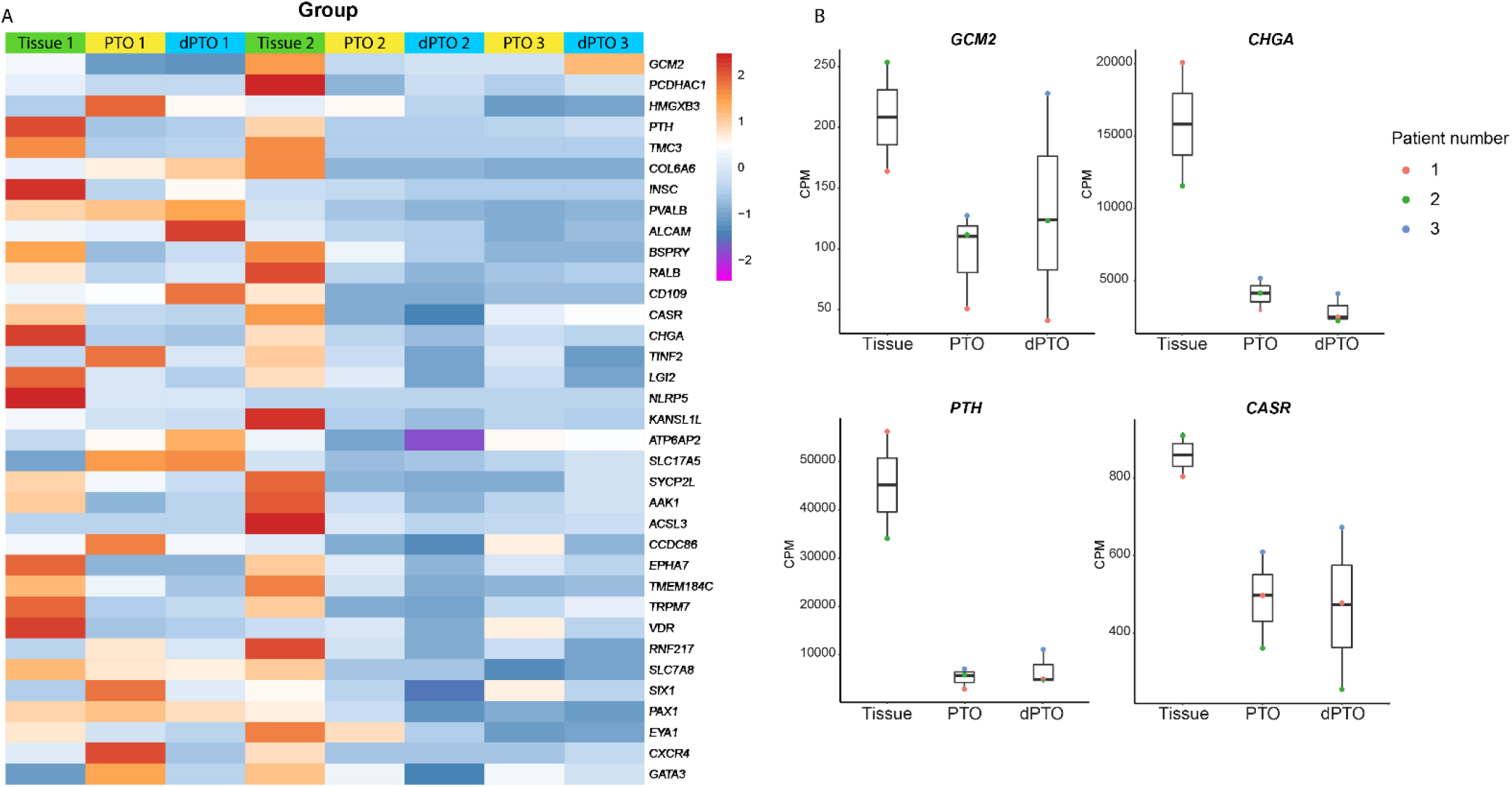
Bulk RNA-sequencing reveals limited differences between primary hyperplastic parathyroid tissue, parathyroid organoids at the end of passage 1 and two weeks differentiated parathyroid organoids. **A.** Heatmap indicating gene expression (counts per million, CPM) of parathyroid-specific genes in primary hyperplastic parathyroid tissue, PTOs at the end of passage 1 and two weeks differentiated PTOs (dPTOs). **B.** Boxplots of parathyroid-specific gene expression (counts per million, CPM) in genes primary hyperplastic parathyroid tissue, PTOs at the end of passage 1 and two weeks dPTOs Boxplot shows median, two hinges (25th and 75th percentile) and two whiskers (largest and smallest value no further than 1.5x inter-quartile range). *PTO=* parathyroid organoids, *dPTO=* differentiated parathyroid organoids.

Genes that were differentially expressed in tissue compared to PTOs and dPTOs (Supplemental Figure 3, Supplemental Table 2 and 3) were categorized and represented as biological processes and pathways (gene ontology (GO) terms, Supplemental Figure 4 and 5, Supplemental Tables 4-7). Interestingly, genes such as *STC1* (Stanniocalcin 1) and *CCL2* (C-C Motif Chemokine Ligand 2) that were significantly upregulated in both PTO and dPTO compared to tissue were amongst others associated with calcium ion transport. (Supplemental Tables 4 and 6). Genes significantly upregulated in the dPTO group compared to tissue seem to involved in the regulation of cell population proliferation (Supplemental Table 6). These organoids were treated following the modified Bingham differentiation protocol, which might be the reason of this possibly more proliferative state, since in PTOs this GO term was not detected (Supplemental Tables 4 and 6). Furthermore, genes significantly upregulated in PTO compared to tissue were associated with angiogenesis and vascularization. No parathyroid-specific biological processes, such as hormone production and calcium ion transport, were associated with downregulated in both PTO and dPTO conditions when compared to tissue (Supplemental Figure 5, Supplemental Tables 5 and 7).

Moreover, we found that all parathyroid-specific genes (31–34) expressed in hyperplastic parathyroid tissue samples were also expressed in PTOs and dPTOs (Figure 3A). *Glial Cells Missing Transcription Factor 2* (*GCM2*), a zinc finger-type transcription factor, is known to be a master regulator for embryonic development of parathyroid glands (35). Since these parathyroid glands are not formed in GCM-null mice, *GCM2* is believed to be essential for development of the parathyroid glands and survival at the earliest stage after organ specification during the embryogenesis (35–38). *Chromogranin A* (*CHGA*) is an acidic protein in secretory granules of (neuro)endocrine cells and is a major protein of the parathyroid gland that is co-stored and co-secreted with PTH (39,40). When we focus on four parathyroid specific genes, i.e., *GCM2, CHGA, PTH* and *CASR*, gene expression of all was observed, albeit to a lower extend, in dPTO and PTO compared to primary tissue (Figure 3B). Immunolabeling of these important parathyroid markers was confirmed at the protein level (Figure 4). When comparing immunolabeling of parathyroid tissue with parathyroid organoids, similarities were seen in the expression pattern of GCM2, CHGA, PTH and CaSR (Figure 4).

**Figure 4.**
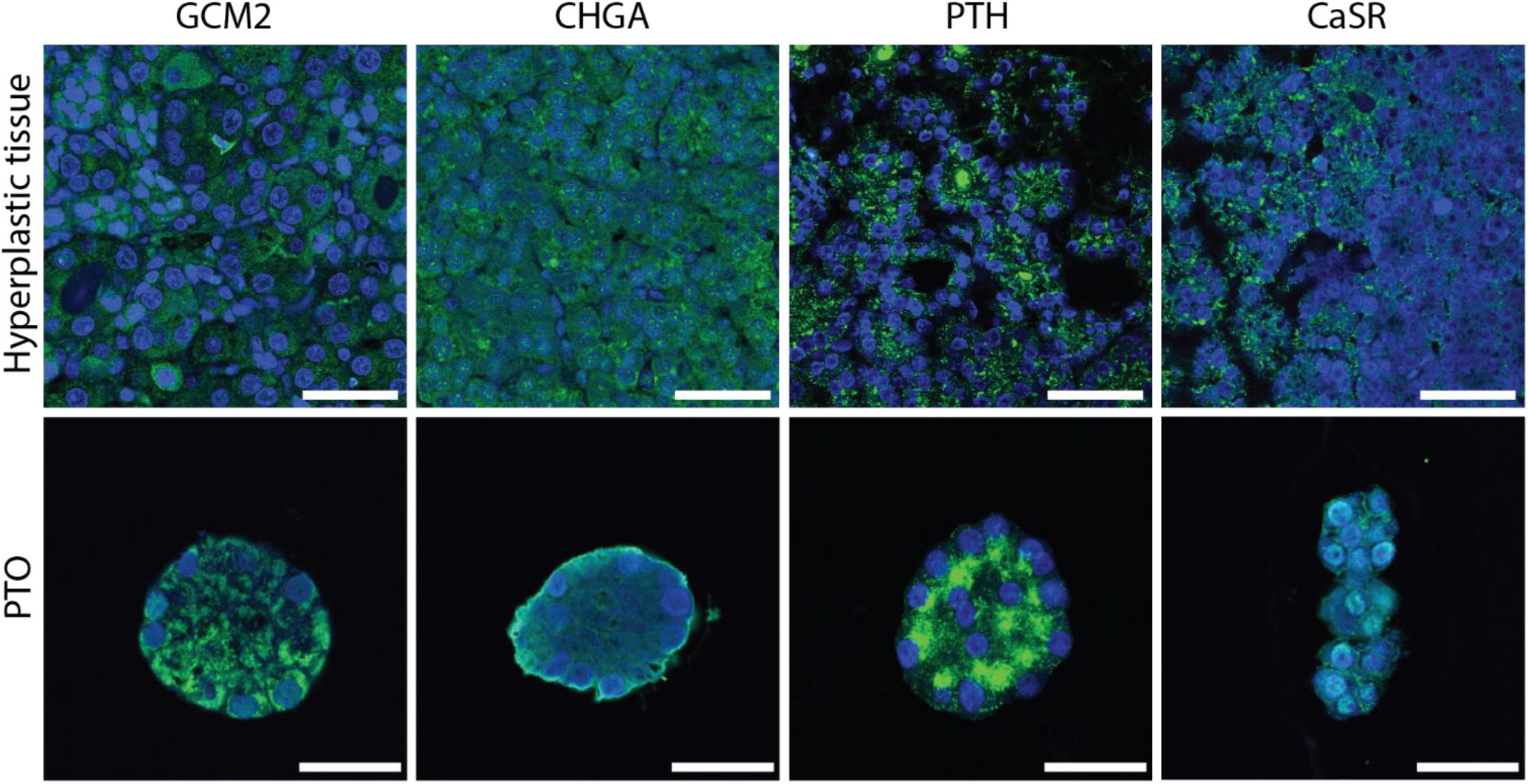
Immunofluorescent characterization of parathyroid organoids. Representative images of organoids and tissue showing parathyroid-specific markers. Scale bar tissue = 100 μm. Scale bar organoids = 25 μm. *PTO=* parathyroid organoid Markers indicated on the top are shown as a green fluorescent signal. Nuclei are shown as a blue fluorescent signal.

To study the presence of stem cell markers in human PTOs, we analyzed the most proposed stem cell markers as listed by van der Vaart *et al* (26), using bulk RNA-sequencing. No upregulation was observed in the putative stem cell markers *MKI67, ATXN1, FUT4, REOX1, HHEX, NGFR, TTF2* and *SOX2*, when comparing PTOs and dPTOs to tissue (Supplemental Figure 6). Additionally, none of the stem cell markers (Supplemental Figure 6) were differentially expressed in both PTOs and dPTOs compared to tissue.

### 2.3 Functionality of parathyroid organoids

PTH secretion is known to be negatively regulated by increased extracellular calcium levels in normal physiology. Therefore, we examined whether the release of PTH by PTO was responsive to extracellular calcium (Figure 5A). We observed a significant decrease of 31.3% ± 10.8% (p=0.015) when organoids were subjected to very high levels of calcium (3.4 mM), when compared to normal culturing conditions (=1.4 mM) (Figure 5B). When the organoids were then switched back to basal 1.4 mM calcium, we observed a significant increase of 53.0% ± 11.1.% (p=0.018) (Figure 5B). These results indicate the fully functional response of PTOs.

**Figure 5.**
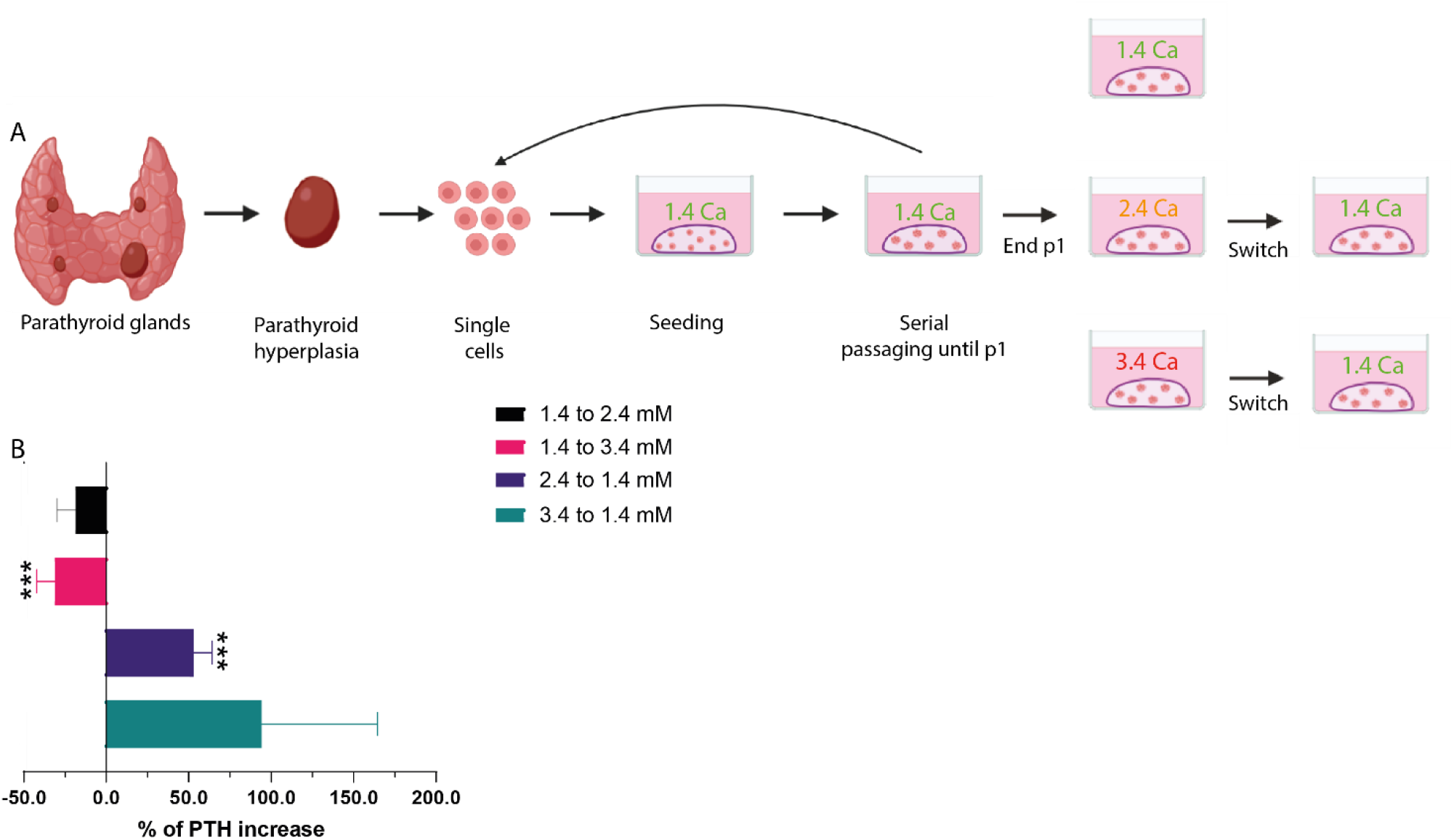
Calcium-response experiment of parathyroid organoids. **A.** Schematic representation of parathyroid tissue isolation, primary tissue culture, and the calcium-response experiment. **B.** Secreted PTH levels from PTOs in switched calcium concentrations (n= 4 patients from 1.4 to 2.4 mM, n= 12 patient from 1.4 to 3.4 mM, n= 4 patients from 2.4 to 1.4 mM, and n=4 patients from 3.4 to1.4 mM, error bars represent the SEM, normalized to PTH levels in the first calcium concentration). ***= significant compared to 1.4 mM (baseline) (p<0.05), *PTO=* parathyroid organoid.

### 2.4 Drug screening

To demonstrate the utility of patient-derived PTO in testing potential novel therapeutic targets, validated therapeutics were used to assess the effect of the drug on PTH secretion by the parathyroid organoids (Figure 6A). Cinacalcet HCl (cinacalcet), an allosteric modulator of CaSR that increases the sensitivity of the CaSR to activation by extracellular calcium, suppresses PTH secretion in patients with primary hyperparathyroidism or secondary hyperparathyroidism on maintenance dialysis (41). Calcitriol, a vitamin D analogue, inhibits the production of PTH by acting directly on the parathyroid gland vitamin D receptor, thereby inhibiting transcription of the gene encoding PTH for the synthesis of PTH and indirectly by increasing calcium uptake in the gastrointestinal system, maintaining PTH secretion at low levels (42,43). We observed a significant decrease in PTH secretion when organoids were subjected to different concentrations of cinacalcet compared to DMSO control (average reduction of 58.9% ±8.5% for 1.5 μM and 58.8% ± 12.9% for 2.5 μM, p=0.020 and p=0.045) (Figure 6B). A similar effect was observed when organoids were exposed to different concentration of calcitriol (average reduction of 50.3% ± 3.3% for 2 nM and 54.4% ± 2.0% for 20 nM, p<0.0001 and p<0.0001, respectively).

**Figure 6.**
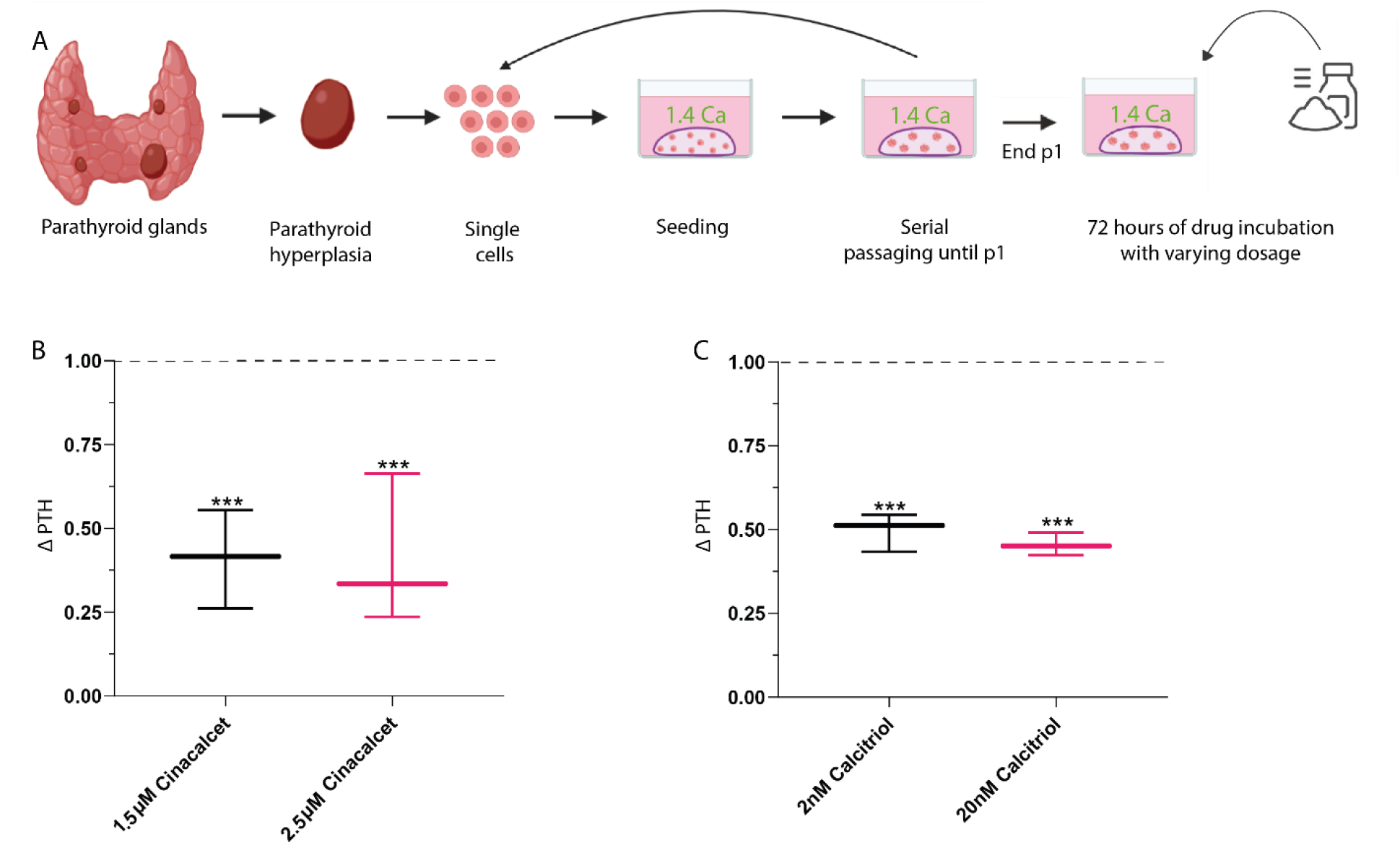
Drug-screening experiment of parathyroid organoids. **A.** Schematic representation of parathyroid tissue isolation, primary tissue culture, and the drug-screening experiment. **B.** Secreted PTH levels from PTOs exposed to 1.5μM and 2.5μM cinacalcet (n=3 patients, error bars represent the SEM, normalized to PTH levels in the DMSO control = dotted line). **C.** Secreted PTH levels from PTOs exposed to 2nM and 20nM calcitriol (n=3 patients, error bars represent the SEM, normalized to PTH levels in the DMSO control = dotted line). ***= significant (p<0.05), *PTO=* parathyroid organoid.

### 2.5 PET/SPECT tracer validation

To demonstrate the utility of patient-derived PTOs to validate PET/SPECT tracers, validated tracers, i.e. ^11^C-methionine (44–47) and ^99m^Tc-sestamibi (48–50), were used to assess uptake of the tracer by the PTOs. ^11^C-methionine is an amino acid tracer used to localize parathyroid adenomas using a PET (positron emission tomography) scan. The ^11^C-methionine uptake mechanism is not yet fully understood; it is presumed to be involved in the synthesis of pre-pro-PTH, a PTH precursor which results in intense and specific uptake in hyperfunctioning parathyroid glands (51,52). Also, the exact mechanism of ^99m^Tc-sestamibi uptake in parathyroid glands remains unknown. ^99m^Tc-sestamibi uptake depends on numerous factors, including perfusion, cell cycle phase and functional activity (53). It has been suggested that the electrical potential of the plasma and mitochondrial membrane regulates uptake of ^99m^Tc-sestamibi and that tissues rich in mitochondria are avid for it (54,55). ^99m^Tc-sestamibi is detected using a gamma-camera which can acquire SPECT (single photon emission computed tomography) 3D images.

We found that the PTOs showed cell-associated binding of both ^11^C-methionine (cell binding in 10,000 organoids 0.14 ± 0.09%) and ^99m^Tc-sestamibi (cell binding in 10,000 organoids 1.05 ± 0.55%) (Figure 7B, D). ^11^C-methionine showed a significant reduction in cell binding when PTOs were blocked with an excess of L-methionine (0.14 ± 0.09% versus −0.14% ± 0.09% p<0.0005), indicating specific binding (Figure 7B, Supplemental Figure 7). Additionally, when PTOs were incubated with ^99m^Tc-sestamibi, we observed a higher uptake in PTOs originating from patients with a ^99m^Tc-sestamibi positive scan as compared to PTOs originating from patients with a ^99m^Tc-sestamibi negative scan (1.05 ± 0.55% versus −0.18% ± 0.07%, p=0.010), possibly indicating that we cultured patient-specific PTOs (Figure 7D, Supplemental Figure 7).

**Figure 7.**
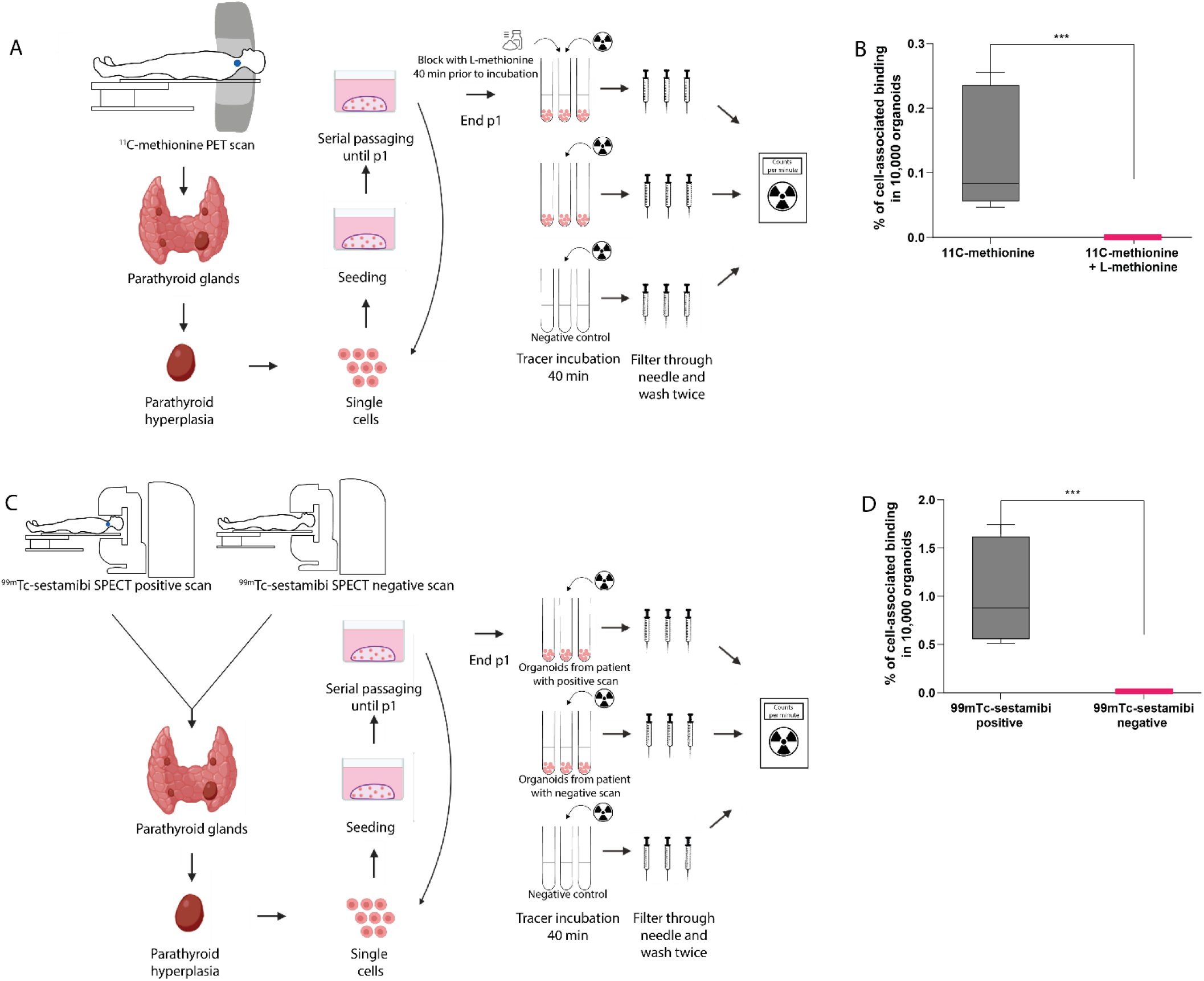
PET/SPECT experiment of parathyroid organoids with existing tracers A, C. Schematic representation of parathyroid tissue isolation, primary tissue culture, and the PET/SPECT experiment. **B.** Percentage of ^11^C-methionine cell-associated binding compared to percentage of cell-associated binding after blocking with L-methionine (n=3 patients, Tukey boxplot). **D.** Percentage of ^99m^Tc-sestamibi cell-associated binding in PTOs originating from patients with a ^99m^Tc-sestamibi positive scan (n=2 patients, Tukey boxplot) compared to percentage of cell-associated binding in PTOs originating from a patient with a ^99m^Tc-sestamibi negative scan (n=1 patient). *PTO*= parathyroid organoid. For visualization purposes, negative uptake values were set at 0%.

## 3. Discussion

In this paper, we report the establishment of a patient-derived parathyroid organoid culture, showing self-renewal potential suggesting the presence of stem cells. Furthermore, we have shown that these PTOs resemble the originating tissue on both gene and protein expression levels and functionality. We illustrate clinically relevant responses (increased and decreased hormone secretion) of these PTOs in response to changes in calcium concentration and PTH lowering drugs. Furthermore, we also found specific parathyroid-targeted tracer uptake in our PTOs, all together demonstrating that these organoids can model human parathyroid functionality.

We confirm that our organoids phenocopy primary parathyroid tissue using RNA-sequencing. In fact, the observed heterogeneity in gene expression between patients seemed to reflect in the respective organoids. The GO biological process of “regulation of cell population proliferation” was significantly upregulated in dPTOs compared to tissue, in contrast to PTOs, suggesting that this upregulation is the result of exposing PTOs to the modified Bingham differentiation protocol. Furthermore, when organoids were exposed to this differentiation protocol for two weeks, it is possible to increase the PTH excretion with an additional 50%.

Regretfully, we were unable to find evidence for the existence of unique parathyroid stem cells in our *in vitro* PTO model. This is in line with results from a recent study by van der Vaart *et al*. in thyroid organoids who could not find evidence for the existence of unique thyroid stem cells (26). They described that many solid tissues do not rely on a professional stem cell for their maintenance and repair, but rather temporarily recruit differentiated cells into a transient progenitor pool (26), suggesting that this phenomenon is also applicable to the thyroid or in this case even parathyroid gland.

For the clinical development of new imaging tracers and drugs, an established appropriate pre-clinical model for testing is required (56). We used our culturing method to culture parathyroid organoids to perform functional testing. Manipulating the calcium concentration and using clinically available PTH-lowering drugs, resulted in a significantly reduced PTH excretion of PTOs, demonstrating that these organoids can model human parathyroid functionality and are a valid *in vitro* model. This effect is similar to the response expected in patients. Next to this, we demonstrated that the PTOs showed specific binding of the PET tracer ^11^C-methionine and SPECT tracer ^99m^Tc-sestamibi. This indicates that we may be able to culture patient-specific PTOs. This methodology could also be used to predict patient response to treatment using our organoid model and develop or test new effective parathyroid imaging tracers.

This study is the first to develop a patient-derived PTO culture methodology of parathyroid tissue. Previous studies aimed at culturing bovine or rat parathyroid glands in 2D (7–12). However, those cell culture models of bovine origin fail due to fibroblast cell overgrowth, and rapid loss of calcium responsiveness (10–12). Since cells *in vivo* experience a complex 3D environment that exposes them to circulating molecules, neighbouring cells, and the extracellular matrix, 2D cell cultures are unable to mimic the complicated microenvironment cells experience in tissue (57). Organoids are self-organizing, 3D culture systems that are highly similar to actual human organs, including increased exposure to circulating molecules, neighbouring cells, and the extracellular matrix (Matrigel) (18,58–62). Organoids from several glandular tissues have been developed, such as thyroid organoids (24), papillary thyroid cancer organoids (63), salivary glands (64), and prostate cancer organoids (65). The analysis of organoid formation could provide valuable insight about the mechanisms underlying human development and organ regeneration (58). Such analysis highlight the value of organoids for discovering potential novel therapeutic targets, opening up opportunities for future imaging tracer testing and targeted treatment screening. We encountered a similar effect in our earlier organoid cultures but lowering the fibroblast growth factor 10 (FGF10) concentration solved this problem. We did see a small increase in fibroblasts in later passages (p3). However, these organoids are significantly larger then in earlier passages (p1/p2), and sometimes lack sufficient support of the Matrigel. Thereby exposing the PTO to the topography of the culture plate potentially leading to epithelial - mesenchymal transition (66). The limitations of our study include the absence of the original microenvironment. This consists of, amongst others, fluctuating concentrations of extracellular signals and the presence of blood vessels. This may be the reason why our PTOs do not have exactly the same transcriptome as hyperplastic parathyroid tissue. The limited differences that were observed, likely do not affect the functionality of our PTOs, since the functional testing and tracer experiment all show that the PTOs are a highly suitable model that resembles functional parathyroid tissue. Furthermore, it has been shown that microenvironmental factors are able to alter the therapy response of cells, complicating the use of organoids for these studies (67,68). Additional research is needed to further explore the clinical application of this parathyroid culture methodology as one day these parathyroid organoids may aid as a stem cell transplantation method for patients with abnormally low levels of PTH (hypoparathyroidism). For this application, first a comparison has to be made between healthy parathyroid tissue and hyperplastic parathyroid tissue.

In conclusion, we present a patient-derived parathyroid organoid culture that recapitulates the originating tissue on gene and protein expression and functionality. This parathyroid culture will be able to provide a novel *in vitro* model to perform physiology studies and discover potential novel therapeutic targets, opening opportunities for future tracer testing and drug screening of the parathyroid gland.

## 4. Materials and Methods

### 4.1 Patient material

Human benign hyperplastic parathyroid tissue was obtained from patients undergoing parathyroid surgery. The biopsies were transported from the operating room (OR) in Hank’s Balanced Salt Solution (HBSS) (Gibco, NY, USA) with 1% bovine serum albumin (BSA) (Gibco), on ice for immediate further processing.

### 4.2 Parathyrosphere cultures

The biopsies were mechanically dispersed using the gentleMACS dissociator (Miltenyi Biotec, Leiden, the Netherlands) and simultaneously subjected to digestion in 5mL HBSS/1% BSA buffer containing 80 mg/mL dispase (Sigma), 1.2mg/mL collagenase type II (Gibco), and calcium chloride (Sigma) at a final concentration of 6.25 mM. Followed by two periods of 15 minutes in a 37°C shaking water bath. This isolate was collected by centrifugation, washed in HBSS/1% BSA solution, and passed through a 100 μm cell strainer (BD Biosciences, NJ, USA). The cell pellet was collected by centrifugation and resuspended in Dulbecco’s modified Eagle’s medium: F12 medium (DMEM:F12) containing Pen/Strep antibiotics (Invitrogen) and Glutamax (Invitrogen). Every 20 μL of this cell solution was combined with 40 μL of Basement Membrane Matrigel (BD Biosciences) on ice and subsequently plated in the center of a 24-well tissue culture plate. Following solidification of the gel for 30 minutes in the incubator, 500 μL of complete growth medium (parathyroid medium) consisting of 50% conditioned Wnt3a medium, 10% conditioned R-Spondin medium and 40% DMEM:F12 containing B27 (Gibco, 0.5x), HEPES (Gibo, 10 mM), ROCK inhibitor Y-27632 (10 μM, Abcam, Cambridge, UK), Nicotinamide (100 uM, Sigma), FGF-10 (50 ng/mL, peprotech), Noggin (25 ng/mL, Peprotech, NJ, USA), EGF (20 ng/mL, Sigma), TGF-β inhibitor A 83-01 (5 uM, Tocris bioscience), VEGF-121 (10 ng/mL, Immunotools) and Angiotensin (0.1 nmol/mL, Sigma) was added and replaced on a weekly basis.

### 4.3 Self-renewal

To study the long-term self-renewing potential, organoids were dissociated and re-plated for the next passage. Prior to passaging, Matrigel was dissolved by incubation with Dispase enzyme (1 mg/mL for 30 minutes at 37 °C; Sigma) and organoids with a size >50 μm were counted using the Ocular Micrometer at a 10x objective. Initial parathyrosphere cultures of seven days old (d) were dissociated to single cells using 0.05% trypsin-EDTA (Invitrogen), counted using Tryphan blue and cell concentration was adjusted to 2 x 10^6^ cells per ml. In total, 20 μL of this cell solution was combined on ice with 40 μL of Basement Membrane Matrigel (BD Biosciences) and plated in the center of a 24-well tissue culture plate. After solidifying the Matrigel for 30 minutes at 37 °C, gels were covered in 500 μL of parathyroid medium as defined above and parathyroid medium was renewed weekly. This self-renewal procedure was repeated every three to four weeks and up to 5 times (4 passages).

### 4.4 Organoid differentiation

PTOs were encapsulated in Matrigel as described above and subjected to a modified Bingham Protocol for differentiation (28,29). Standard culture medium, DMEM:F12 containing Pen/Strep antibiotics (Invitrogen), and Glutamax (Invitrogen). In short, cells were continuously exposed to 100 ng/mL Activin A (Immunotools), 100 ng/mL Shh (R&D systems) and 5% FBS for 14 days with media changes every seven days.

### 4.5 PTH secretion

Medium was collected before passage and before each time point during the differentiation period, and stored at −20 °C until use. Medium was used to measure the concentration of secreted PTH protein by the PTOs. Secreted PTH was measured by electro-chemical luminescence immuno assay (ECLIA) using an Elecsys PTH kit (Roche, Germany).

### 4.6 Quantitative Polymerase Chain Reaction

Total RNA from patient material (n=3) and organoids (p1 and p2 n=3, p3 n=2) was extracted (RNeasy™ Mini Kit, Qiagen). To obtain cDNA, 500ng of total RNA was reverse transcribed using 1 μL 10 mM dNTP Mix, 100 ng random primers, 5x First-strand Buffer, 0.1 M DTT, 40 units of RNase OUT and 200 units of M-MLV RT, in a volume of 20 μL for each reaction (all Invitrogen). qPCR (Bio-Rad) was performed using Bio-Rad iQ SYBR Green Supermix according to manufacturer’s instructions. For each sample, PCR buffer was mixed with 100 ng cDNA, sybergreen and forward and reverse primers for the targeted genes in a volume of 13 μL. A three-step qPCR reaction was applied. Oligo sequences of primes used were as followed: PTH fwd, 5’-AGCTACTAACATACCTGAACG-3’; PTH rev, 5’-CTCTCCATCGACTTCAGATG-3’; CaSR fwd, 5’-AGATGGCACGGGACACTACC-3’; CaSR rev, 5’-AGGAGGCATAACTGACCTGGG-3’; GATA3 fwd, 5’-AAGCCTCTGCAATGTGCTC-3’; GATA3 rev, 5’-GTGGTGGTCTGACAGTTCGC-3’, and YWHAZ fwd, 5’-GATCCCCAATGCTTCACAAG-3’; YWHAZ rev, 5’-TGCTTGTTGTGACTGATCGAC −3’.

### 4.7 RNA bulk sequencing

Total RNA was extracted from patient material (n=3), organoids at the end of passage one (n=3) and organoids at the end of passage one subjected to two weeks of differentiation (n=3) using the RNeasy Mini Kit (Qiagen, 74104) according to the manufacturer’s protocol. RNA quality and quantity were determined using High Sensitivity RNA ScreenTapes (Agilent, 5067). Eight RNA samples, with RIN values varying between 7.1 and 9.2 were used for library preparation according to the manufacturer’s protocol. For one of the tissue samples, no RNA quality could be detected using the High Sensitivity RNA ScreenTape, therefore we decided to exclude this sample for further analysis. Quality and concentration of libraries from individual samples were assessed using the Fragment Analyzer (Agilent). Subsequently, individual libraries were combined into a sequencing pool of 8 samples each with equal molar input. 2 pM were loaded, and 75 bp paired-end sequencing was performed on an Illumina NextSeq 500 system for a 75 bp paired-end sequencing run.

### 4.8 RNA sequencing analysis

All bioinformatic analyses were performed with available packages in RStudio (v4.0.2) and plots were generated with ggplot2 (v3.3.3.9000) (69). Heatmaps were generated with pheatmap (v1.0.12) (70), and row clustering was performed with the ward.D2 clustering method. For the differential gene expression analysis, lowly expressed genes (total count < 1 in less than 2 samples) were excluded. The bioconductor package edgeR (v3.32.1) (71) was used for normalization by using trimmed mean of M-values (TMM) and identification of differential expressed genes (logFC </>0.95, FDR<0.05) by fitting a generalized linear model.

Gene ontology (GO) analysis for differentially expressed genes was performed with enrichR (v 3.0) (72). The database “GO_Biological_Process_2021” was used to identify enriched GO terms.

### 4.9 Immunostaining

For immunostaining, antibodies to PTH (1:100, Boster), CaSR (1:100, abcam), GCM2 (1:100, proteintech) and CHGA (1:100, proteintech) were used to detect proteins. In the case of the organoids, Matrigel was dissolved by incubation with Dispase enzyme. Followed by subsequent washes with PBS/0.2% BSA and centrifuged at 400 g for 5 min. The resulting pellet was fixed in 4% paraformaldehyde (15 min, RT) and washed with PBS. Next, the organoids were embedded in HistoGel (Richard-Allan Scientific/Thermo scientific) and the gel was subjected to dehydration, followed by embedding in paraffin and sectioning (5 μm). Patient material was fixed in 4% formaldehyde. After dehydration, the tissue was paraffin-embedded and sectioned at a 5 μm thickness. Sections were de-paraffinized and subsequently, Tris-EDTA antigen-retrieval was performed. This was followed by washing, blocking, and incubation with primary antibodies overnight. Slides were then incubated with secondary antibodies, stained with 4’,6-diamidino-2-phenylindole (DAPI), mounted using aqueous mounting medium (DAKO), and imaged using a DM6B microscope (Leica) and a TCS SP8 X confocal microscope (Leica).

### 4.10 Effect of extracellular calcium on PTH secretion

Undifferentiated organoids were exposed to parathyroid medium containing normal (1.4 mM), high (2.4 mM) or higher (3.4 mM) concentration of calcium for 7 days (30,73). Medium calcium content was measured by photometrical absorption measurement of calcium EDTA complex using a Calcium Gen.2 kit (Roche, Germany). After 7 days, the medium was collected to measure the secreted concentration of PTH. Hereafter, organoids were switched from normal (1.4 mM) to higher (3.4 mM) concentration of calcium or vice versa to study their rescue potential. After 7 days, the medium was again collected to measure the secreted concentration of PTH.

### 4.11 Drug screening

Cinacalcet HCL (Combi-Blocks, QA-8513) was added to the medium in two different concentrations (1.5 μM and 2.5 μM) and following 72 hours of incubation, the medium was collected to measure the concentration of secreted PTH protein by the PTOs. The concentration of Cinacalcet HCL used was based on literature that investigated the effect on parathyroid cultured cells (74,75). Further, calcitriol (Cayman Chemical, 32222-06-3) was added to the medium in two concentrations (2 nM and 20 nM) and following 72 hours of incubation, the medium was collected to measure the concentration of secreted PTH protein by the PTOs. The concentration of calcitriol used was also based on a previous study (76). Both cinacalcet and calcitriol were dissolved using DMSO, therefore DMSO was used as vehicle control in a concentration of 0.1%.

### 4.12 PET/SPECT tracer validation

To demonstrate the utility of patient-derived PTOs to validate PET/SPECT tracers, existing tracers, i.e. ^11^C-methionine and ^99m^Tc-sestamibi, were used to assess uptake of the tracer by the PTOs.

Prior to the experiments, the Matrigel was dissolved by incubation with Dispase enzyme followed by three washes with 1000 μL of HBSS/1% BSA. Approximately 10,000 organoids in 450 μL HBSS/1% BSA were incubated with 2 MBq (in 50 μL) tracer solution at 37 °C for 40 minutes. For ^11^C-mehionine, organoids were blocked with 2 μM (in 50 μL) of cold L-methionine (Sigma-Aldrich) 40 minutes prior to tracer incubation. After tracer incubation, the organoids were collected with a 5-micron filter needle and subsequently washed twice through the filter needle with 500 μL ice-cold HBSS/1% BSA. Activity in the organoids was measured in a gamma-counter (Wizard^2^ 2480, SW 2.1, PerkinElmer). Tracer uptake was corrected for the number of organoids by counting the number of organoids organoids using the Ocular Micrometer at a 10x objective prior to the experiment. Uptake was expressed as percentage of organoid-associated radioactivity per 10,000 organoids. Uptake values were corrected for background radiation. All experiments were executed in duplicate or triplicate, depending on the number of organoids.

### 4.13 Data availability

The raw bulk RNA-Seq data (fastq files) used in this study have been deposited in the NCBI’s Gene Expression Omnibus (GEO) (accession number GSE197445).

### 4.14 Statistics

All data represent the mean ± SEM. Statistical analyses and producing graphs were done using GraphPad Prism, version 8 (GraphPad Software). The statistical methods relevant to each figure are described in the figure legends. Statistical comparisons between two groups were performed using the Mann-Whitney U test or Student’s t test (depending on normal distribution). P values of less than 0.05 were considered to indicate a significant difference between groups.

### 4.15 Study approval

Written informed consent of participants was obtained prior to participation. The study was approved by the Medical Ethics Committee Groningen (approval no. 2017/640).

## Supporting information

Supplemental tables sequencing

## 5. Author contributions

Conceptualization, M.E.N., L.H.J.S., R.P.C., and S.K.; methodology, M.E.N., and L.H.J.S.; formal analysis, M.E.N., L.H.J.S., L.K., I.F.A., R.W.; investigation, M.E.N., I.F.A. and L.H.J.S.; resources, R.P.C., I.F.A., W.S., A.H.B, W.K, L.J., W.T.Z., A.K. and S.K.; data curation, M.E.N., L.H.J.S., R.P.C., and S.K.; writing—original draft preparation, M.E.N., and L.H.J.S.; writing—review and editing, M.E.N., L.H.J.S., L.K., I.F.A., R.W., W.K., A.K., W.S, W.T.Z., L.J., A.H.B., R.P.C. and S.K.; visualization, M.E.N., and L.H.J.S.; supervision, R.P.C. A.H.B. and S.K.; project administration, R.P.C. and S.K.; funding acquisition, R.P.C., A.H.B. and S.K.

All authors have read and agree to the published version of the manuscript.

## 6. Acknowledgements

We would like to thank the surgeons and pathology departments of the University Medical Center Groningen, Treant location Bethesda, and the Martini Hospital for collecting and distributing hyperplastic parathyroid gland tissue. Sequencing was performed with the excellent assistance from Diana Spierings and the ERIBA Research Sequencing Facility. Part of the work was performed at the UMCG Imaging and Microscopy Center (UMIC).

## 8. Supplemental Figures

**Supplemental Figure 1.**
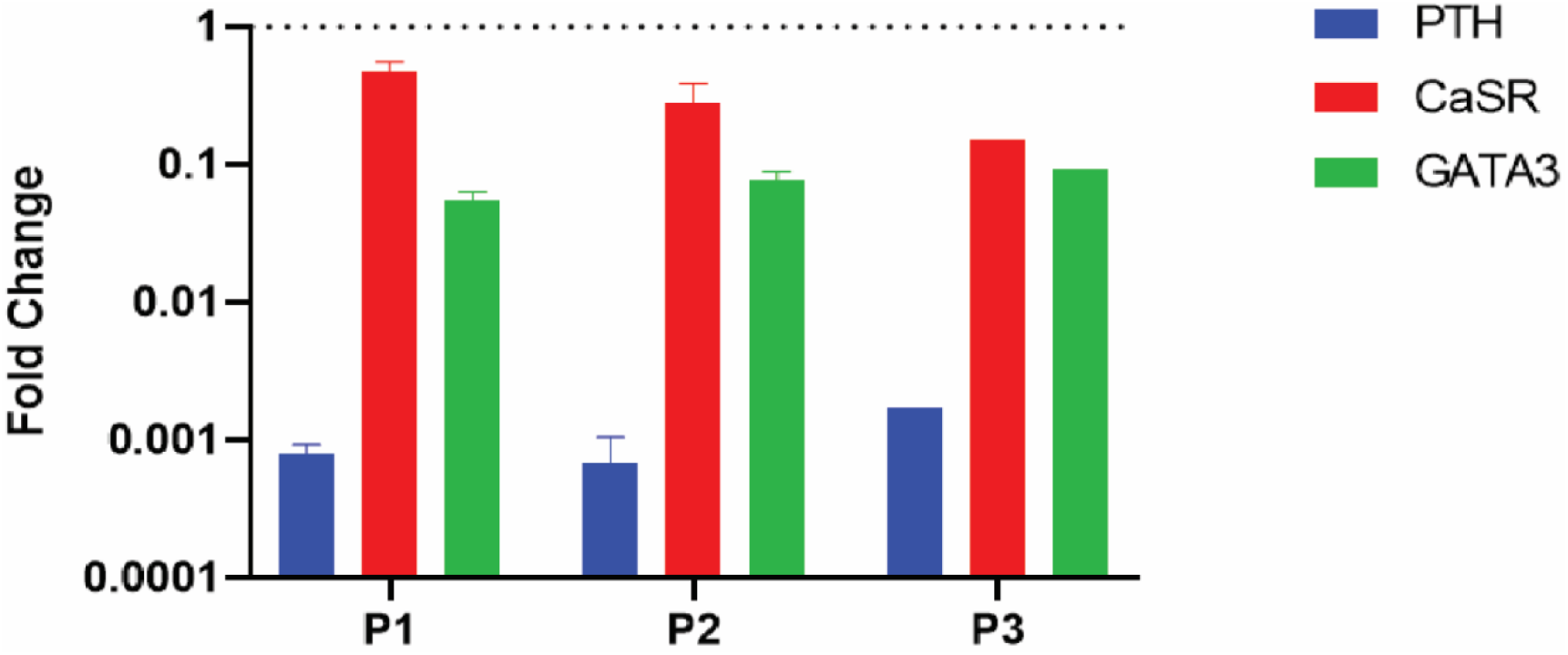
Gene expression pattern analysis across multiple passages (n=3 patients for p1 and p2, n=2 for p3). Dotted line resembles tissue expression levels (n=3), and error bars resemble SEM. No significance was observed between passages with the lowest p-value being 0.065.

**Supplemental Figure 2.**
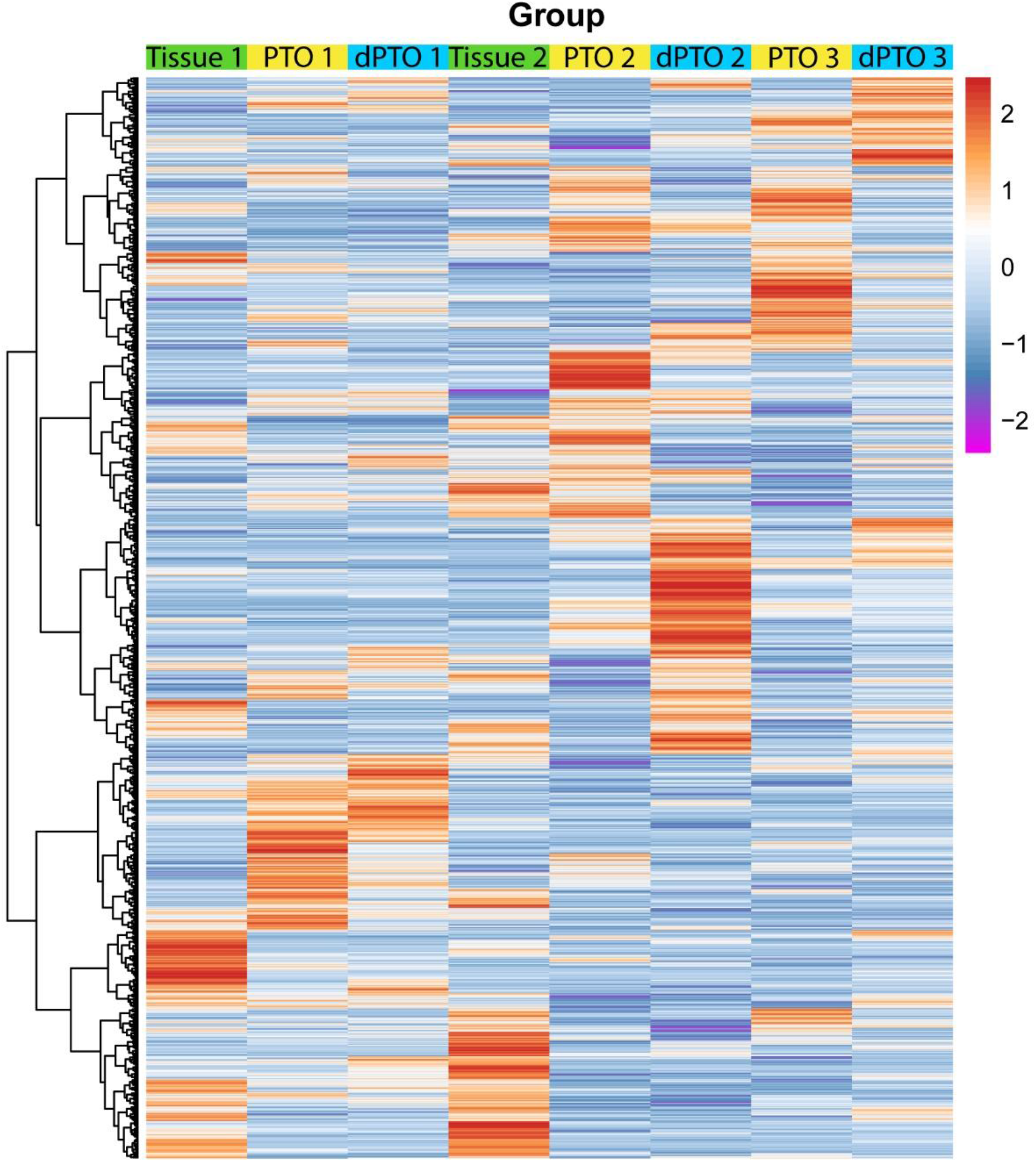
Heatmap showing all sequenced genes for all included samples separately. Data is shown in row Z-score of counts per million (Supplemental table 1). *PTO*= parathyroid organoids, *dPTO*= differentiated parathyroid organoids.

**Supplemental Figure 3.**
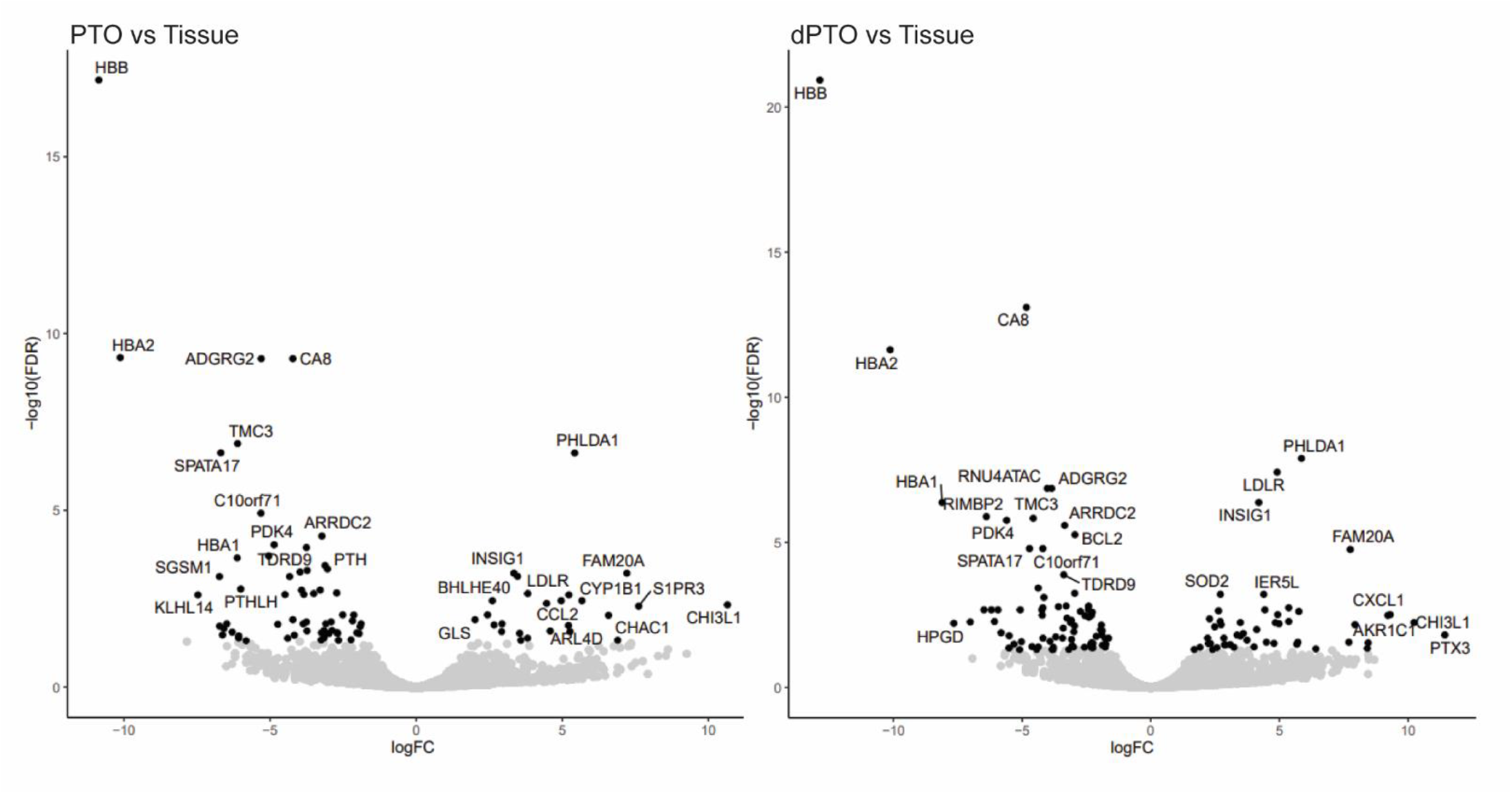
Volcano plots showing differentially expressed genes (FDR<0.05) in PTO or dPTO compared to tissue (Supplemental Table 2 and 3). *PTO*= parathyroid organoids, *dPTO*=differentiated parathyroid organoids.

**Supplemental Figure 4.**
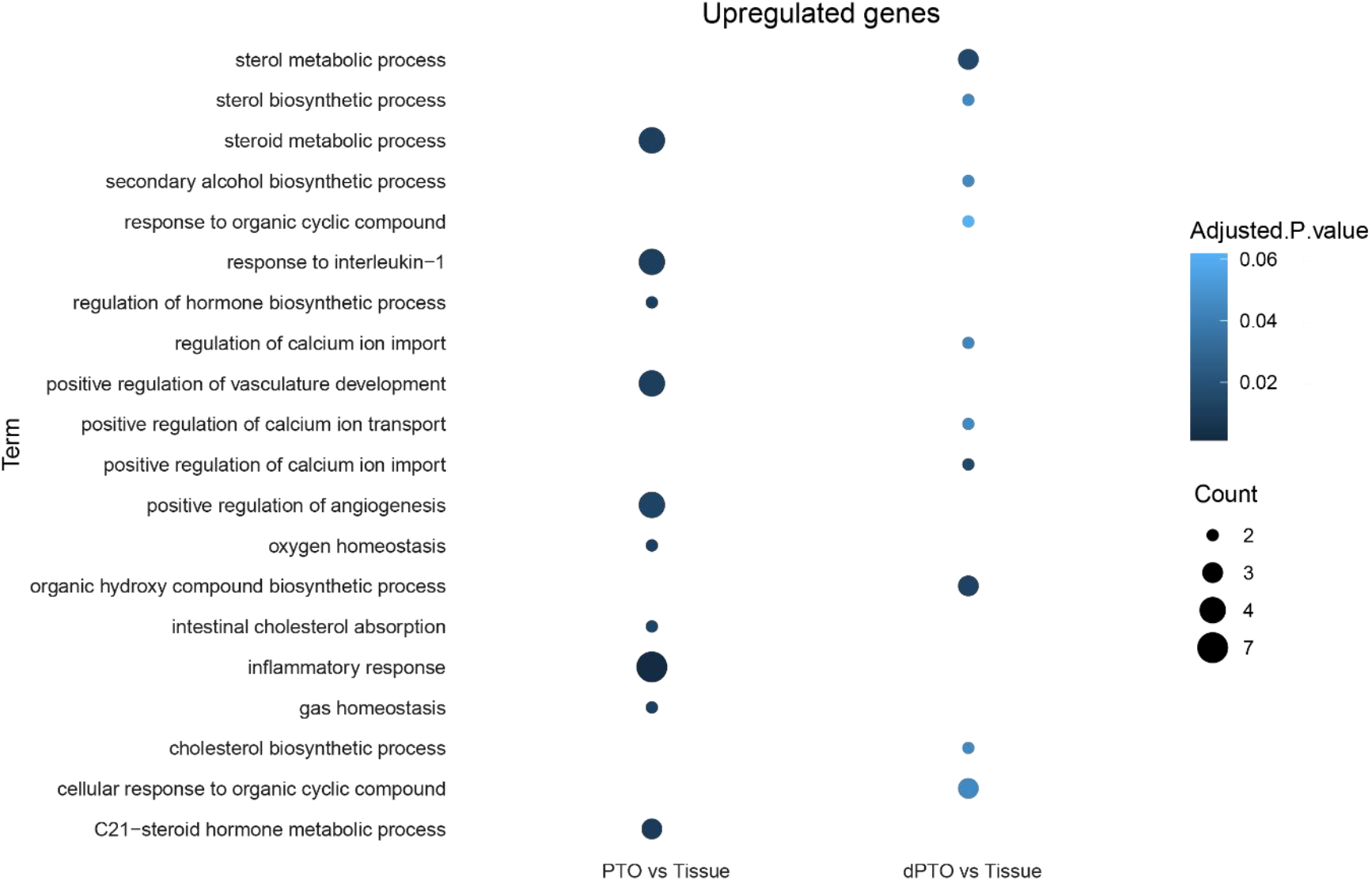
Dotplots showing top ten GO terms associated with upregulated genes in PTO and dPTO compared to tissue. Ranking of GO terms was based on the most significant GO terms. Dot colors resemble significance and size resembles number of associated genes *PTO*= parathyroid organoids, *dPTO*= differentiated parathyroid organoids.

**Supplemental Figure 5.**
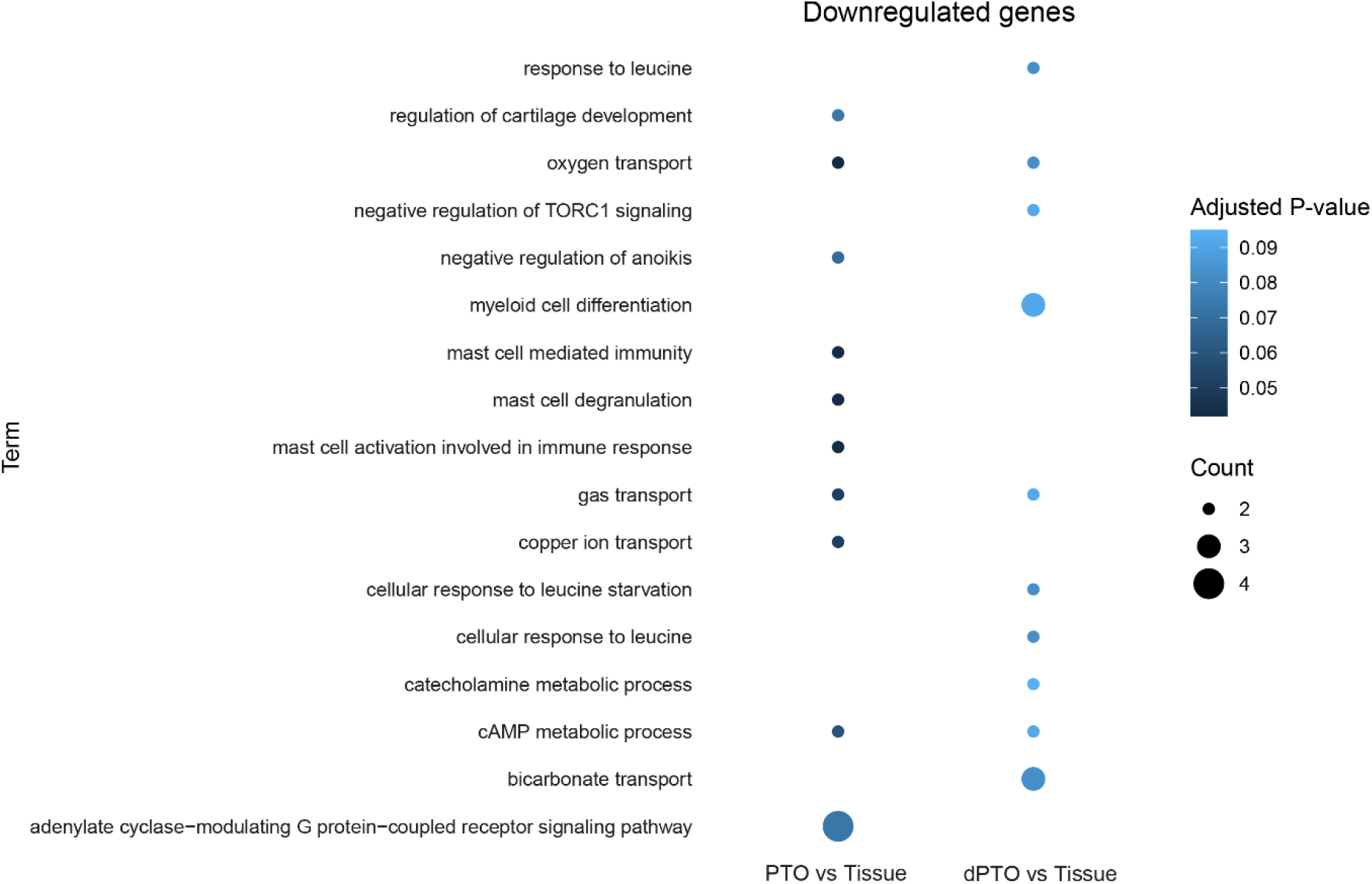
Dotplots showing top ten GO terms associated with downregulated genes in PTO and dPTO compared to tissue. Ranking of GO terms was based on the most significant GO terms. Dot colors resemble significance and size resembles number of associated genes *PTO*= parathyroid organoids, *dPTO*= differentiated parathyroid organoids.

**Supplemental Figure 6.**
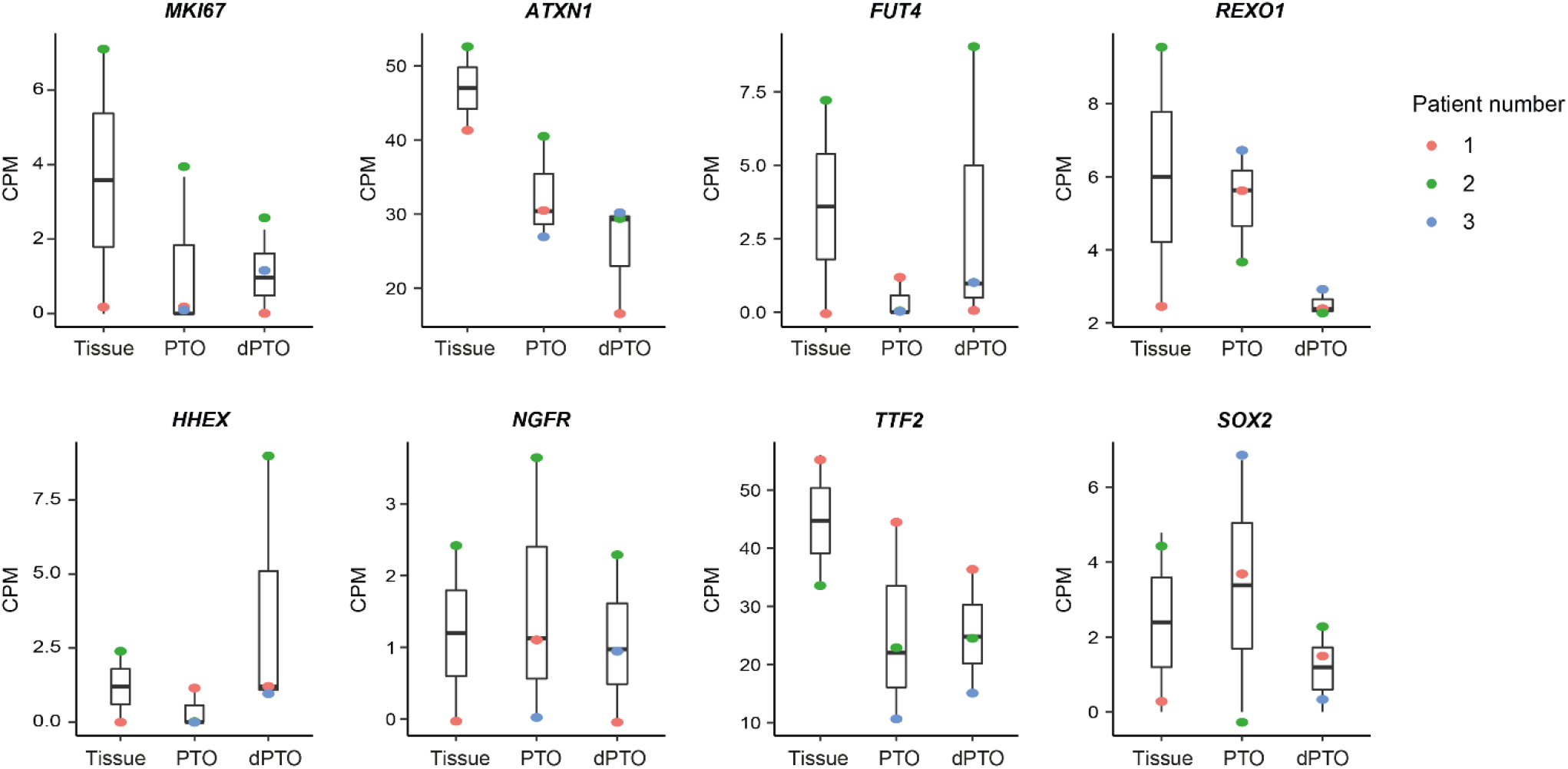
Boxplots of generic stem cell markers of primary hyperplastic parathyroid tissue, PTOs at the end of passage 1 and two weeks dPTOs in counts per million (CPM). Boxplot shows median, two hinges (25th and 75th percentile) and two whiskers (largest and smallest value no further than 1.5x inter-quartile range). *PTO*= parathyroid organoids, *dPTO*= differentiated parathyroid organoids.

**Supplemental Figure 7.**
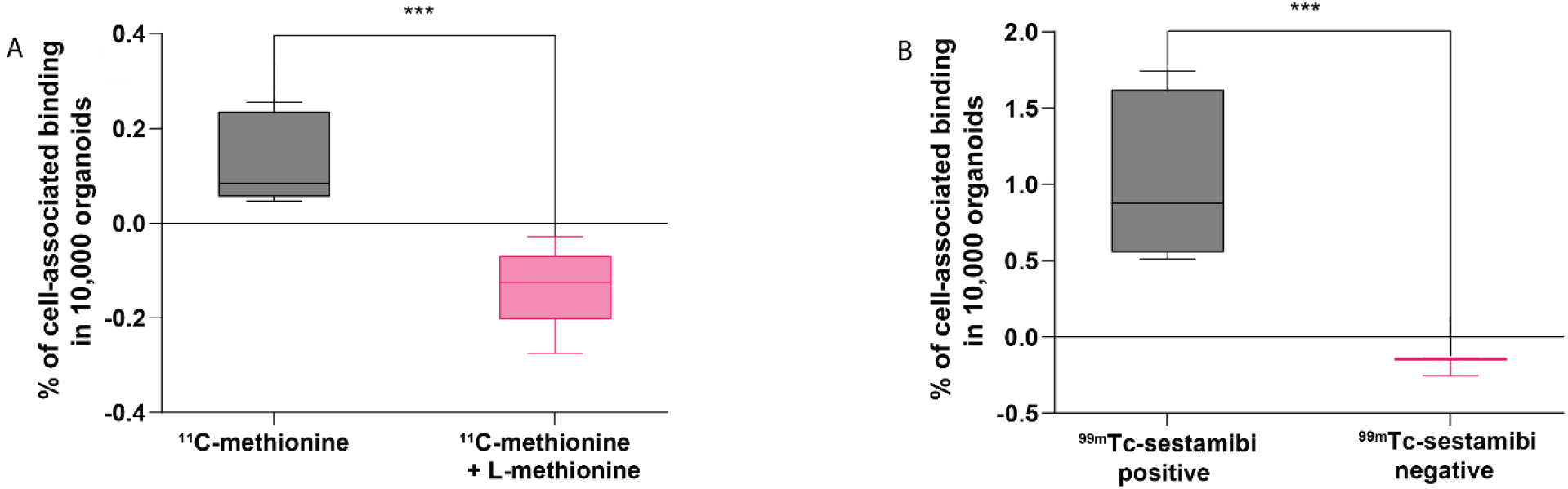
PET/SPECT experiment of parathyroid organoids with existing tracers **A.** Percentage of ^11^C-methionine cell-associated binding compared to percentage of cell-associated binding after blocking with L-methionine (n=3 patients, Tukey boxplot). **B.** Percentage of ^99m^Tc-sestamibi cell-associated binding in PTOs originating from patients with a ^99m^Tc-sestamibi positive scan (n=2 patients, Tukey boxplot) compared to percentage of cell-associated binding in PTOs originating from a patient with a ^99m^Tc-sestamibi negative scan (n=1 patient). *PTO*= parathyroid organoid.

